# Mouse Memory CD8 T cell subsets defined by Tissue Resident Memory (T_RM_) Integrin Expression Exhibit Distinct Metabolic Profiles

**DOI:** 10.1101/2022.09.21.508875

**Authors:** Mike Sportiello, Alexis Poindexter, Emma C. Reilly, Adam Geber, Kris Lambert Emo, Taylor N. Jones, David J. Topham

## Abstract

Tissue-resident memory CD8 T cells (T_RM_) principally reside in peripheral non-lymphoid tissues such as lung and skin and confer protection against a variety of illnesses ranging from infections to cancers. The functions of different memory CD8 T cell subsets have been linked with distinct metabolic pathways and differ from other CD8 T cell subsets. For example, skin-derived memory T cells undergo fatty acid oxidation and oxidative phosphorylation to a greater degree than circulating memory and naïve cells. Lung T_RM_ cells defined by the cell surface expression of integrins exist as distinct subsets that differ in gene expression and function. We hypothesize that T_RM_ subsets with different integrin profiles will utilize unique metabolic programs. To test this, differential expression and pathway analysis were conducted on RNAseq datasets from mouse lung T_RM_ yielding significant differences related to metabolism. Next, metabolic models were constructed and the predictions were interrogated using functional metabolite uptake assays. The levels of oxidative phosphorylation, mitochondrial mass, and neutral lipids were measured. Furthermore, to investigate the potential relationships to T_RM_ development, T-cell differentiation studies were conducted *in vitro* with varying concentrations of metabolites. These demonstrated that lipid conditions impact T cell survival, and that glucose concentration impacts the expression of canonical T_RM_ marker CD49a, with no effect on central memory-like T-cell marker CCR7. In summary, it is demonstrated that mouse resident memory T cells subsets defined by integrin-expression in the lung have unique metabolic profiles and that nutrient abundance can alter differentiation.

## 1 Introduction

Tissue-resident memory T cells (T_RM_) and tumor-infiltrating lymphocytes (TIL) with a T_RM_-like phenotype (CD103+ CD49a+) confer protection against a variety of illnesses ranging from infectious diseases to cancers. As demonstrated in the skin, transfer of T_RM_ from immune to naïve hosts is sufficient to prevent herpes simplex virus type 1 (HSV-1)-associated pathology and, comparably, the presence of antigen-specific CD49a+ CD103+ TIL in a mouse model of melanoma can limit tumor growth (1, 2). In the lungs, T_RM_ are critical for protection against SARS-CoV-2 (COVID-19), influenza, and respiratory syncytial virus, even in the absence of antibodies specific for those viruses (3, 4, 5, 6). T_RM_ have been shown to act as both antigen-specific sentinels linking the innate and adaptive immune systems as well as cytotoxic mediators (7). This functional capacity makes them important players in immunity across multiple organ systems, particularly barrier tissues where nutrient substrate availability varies, and disease states.

T_RM_ are heterogeneous in terms of integrin expression, migration, effector potential, and gene expression (7). Furthermore, the functions of different memory CD8 T cell subsets have been linked with distinct metabolic pathways. For example, skin T_RM_ undergo fatty acid oxidation to a greater degree than other circulating memory and naive cells which tend to have higher glycolytic fluxes (8, 9). However, it is unclear if this programming is universal among all memory T-cell subsets and whether it holds true in other tissues such as the lung. It is also not yet known whether T_RM_ CD8 T cell subsets that express different combinations of surface integrins CD49a and CD103 share metabolic profiles. RNAseq on these populations revealed distinct metabolic transcriptomes (7), suggesting the overall metabolic profiles may differ, which was supported by ex vivo assays. *In vitro* studies to drive differentiation into T_RM_-like phenotypes were used to test whether manipulating nutrient availability affected integrin-based subset development.

## 2 Materials and Methods

### 2.1 RNA sequencing analysis

RNA sequencing data and lists of differentially expressed genes were obtained from previously published work from our lab (GSE179653). Full code that generated those differentially expressed genes as well as further analyses and figures is publicly available at https://github.com/sportiellomike/Lung-CD8-T-cell-Immunometabolism.

### 2.2 Metabolic modelling

Mouse metabolic model iMM1865 was used as a starting point, onto which our publicly accessible transcriptomic data for CD49a^pos^CD103^pos^ (<u>D</u>ouble <u>P</u>ositive, DP), CD49a^pos^CD103^neg^ (<u>CD49a</u> <u>S</u>ingle <u>P</u>ositive, CD49a SP), or CD49a^neg^CD103^neg^ (<u>D</u>ouble <u>N</u>egative, DN) was applied using the Gene Inactivity Moderated by Metabolism and Expression (GIMME algorithm) (10). A threshold value of the median transcript per kilobase million (TPM) for the 1,865 genes in the metabolic model was used. Reaction constraints were added according to previously published research (11). All code used to generate input data, figures, and the models themselves is available at https://github.com/sportiellomike/Lung-CD8-T-cell-Immunometabolism.

### 2.3 Mice

Mice were housed in approved microisolator cages, within a pathogen-free vivarium facility that is accredited by the Association for Assessment and Accreditation of Laboratory Animal Care. The vivarium is staffed with a number of personnel including those with expertise in husbandry, technical skills, and veterinary personnel. C57BL/6J mice (Jackson Laboratories) used for experiments were infected 8-10 weeks after birth. This study was carried out in strict accordance with the recommendations in the *Guide for the Care and Use of Laboratory Animals* as defined by the NIH. The Institutional Animal Care and Use Committee of the University of Rochester approved all protocols.

### 2.4 Virus and infection

Mice were anesthetized with 3,3,3-tribromoethanol. Upon verification of sedation, mice were infected intranasally with 10^5^ EID_50_ of HKx31 human influenza A virus in 30uL volume. After observation to ensure they recovered from anesthesia, mice were monitored daily for weight and overall morbidity to comply with all regulations.

### 2.5 Lung and Bronchoalveolar lavage processing

Bronchoalveolar lavage (BAL) was collected by inserting a flexible Teflon catheter into a slit in the trachea in proximity to the larynx. A 1mL syringe was connected to the catheter and the lungs were flushed 3 times with 1x PBS (volumes of 1mL, 900uL, and 800uL). BAL was spun down at 424xg for 10 minutes. Red blood cells were eliminated through ACK mediated lysis. Cells were washed in 1x PBS containing 1% FBS (PBS+serum) and resuspended in PBS+serum.

The lungs were harvested and separated into right and left lobes prior to collection into RPMI + 8% FBS (Corning #10040CV). Lungs were transferred into Miltenyi Gentle MACS tubes containing 2 mL of 2mg/mL collagenase II (Worthington #LS004177) and mechanically ground up by a Gentle MACS machine. 3 additional mL of 2 mg/mL collagenase II solution was added before incubation for 30 min at 37° C. Further mechanical digestion by Gentle MACS was done at this point. Tubes were spun at 233xg at room temperature for 2 minutes.

Pellets were then dissolved in existing supernatant and transferred through 100 µm strainers. Contents were quantitatively transferred using 5 mL of RPMI + 8% FBS. Tubes were spun at 424xg for 6 minutes at room temperature. The supernatant was discarded, and dissolved in 4 mL 40% Percoll solution, which was then layered on 3 mL 75% Percoll solution. These tubes were then spun at 931xg for 20 minutes at room temperature with no centrifuge break so as not to disturb the layers of the gradient.

The top layer of lipid-rich liquid was aspirated off, and the interface (approximately 2 mL) of the two Percoll densities was collected where the cells were. Tubes were filled with RPMI + 8% FBS and spun down at 424xg for 6 minutes at room temperature.

### 2.6 Spleen processing to achieve single-cell suspension

C57BL/6J mouse spleens were obtained from sacrificed mice and mechanically digested by grinding between two frosted microscope slides without exogenous enzyme added. After grinding, Single-cell suspensions were passed through a plastic mesh to capture large non-single cellular material and centrifuged in conical vials at 424xg at 4°C for 6 minutes. Cells were resuspended in 3 mL of ACK lysis buffer (Cold Spring Harbor Protocol: doi:10.1101/pdb.rec083295) for 5 minutes at room temperature. Volume was brought up to 15 mL in PBS + serum followed by centrifuging at 424xg at 4°C for 6 minutes.

### 2.7 T_RM_-like Media preparation

The following were added to 500 mL DMEM (Corning Catalog No. 10-017-CV): 50 mL fetal bovine serum, 5 mL L-glutamine (Fisher Scientific Catalog No. 25030081), 12.5 mL HEPES buffer (Fisher Scientific Catalog No. 15630080), 6.5 MEM Non-Essential Amino Acids Solution (Fisher Scientific 11140050), 0.6 mL of 1:300 dilution of beta-mercaptoethanol in PBS, 5 mL penicillin-streptomycin (Fisher Scientific Catalog No. 15140122), 0.6 mL gentamycin (Fisher Scientific Catalog No. 15710064). Contents were sterile-filtered.

### 2.8 Fatty Acid Uptake

Fatty Acid Uptake Media (FUM) was created by dissolving 2 µL stock solution made according to package insert per 1 mL of Assay Buffer (AAT Bioquest Screen Quest Fluorimetric Fatty Acid Uptake Assay Kit #36385). Once a single cell suspension was obtained from the harvesting protocol, cells were plated in 100 µL of FUM and 100 µL Leibowitz media for a final volume of 200 µL. Vehicle control used equivalent volume of dimethyl sulfoxide (DMSO) in Assay Buffer. Cells were incubated for 30 minutes at 37° C, then washed 1.5 times with PBS + 1% serum before staining.

### 2.9 Neutral Lipid Staining

Bodipy 505/515 (Fisher Scientific Catalog No. D3921) was dissolved in DMSO to yield a stock of 1 mg/µL concentration. Once a single cell suspension was obtained from the harvesting protocol, each sample of cells in 1 mL of PBS were inoculated with Bodipy to yield a final concentration of 2 mg/mL, or DMSO for the vehicle control. Samples were incubated for 30 minutes at 37° C, then washed 1.5 times with PBS + 1% serum before staining.

### 2.10 2-(N-(7-Nitrobenz-2-oxa-1,3-diazol-4-yl)Amino)-2-Deoxyglucose Uptake

2-(N-(7-Nitrobenz-2-oxa-1,3-diazol-4-yl)Amino)-2-Deoxyglucose (2-NBDG) (Abcam Catalog No. ab146200) was dissolved in DMSO to yield a 20 mM stock solution. After a single cell suspension was obtained from the harvesting protocol, cells were resuspended in 500 µL non-supplemented Leibowitz media and incubated at 37° C with a final 2-NBDG concentration of 80 µM or DMSO for the vehicle control for 35 minutes at 37° C. Samples were washed 1.5 times with PBS + 1% serum before staining.

### 2.11 Mitochondrial Mass Quantification

Mitotracker Green FM (Mitotracker) (Fisher Scientific Catalog No. M7514) was dissolved in DMSO to obtain a 1 mM stock solution. After a single cell suspension was obtained from the harvesting protocol, each sample was dissolved in 500 µL PBS + 1% serum. Samples were inoculated with Mitotracker for a final concentration of 50 nM or equivalent volume of DMSO for the vehicle control and incubated for 45 minutes at 37° C.

### 2.12 Oxidative Phosphorylation Quantification

Tetramethylrhodamine, Ethyl Ester, Perchlorate (TMRE) (Cayman Chemical Catalog No. 701310) was prepared in a 0.5 mM working stock by dissolving in DMSO, according to package insert. After a single cell suspension was obtained from the harvesting protocol, TMRE was added to two separate aliquots of 5 mL RPMI + 8% FBS (TMRE Media) for a final concentration of 250 nM. To one of these aliquots, 2-[2-[4-(trifluoromethoxy)phenyl]hydrazinylidene]-propanedinitrile (FCCP) was added (FCCP-TMRE Media) for a final concentration of 1 µM. Samples were split into two and dissolved into 500 µL of either TMRE Media or FCCP-TMRE Media. Samples were incubated at 37° C for 30 minutes. For vehicle control, DMSO was added in place of TMRE. Cells were washed 1.5 times with PBS + 1% FBS prior to staining.

### 2.13 T_RM_-like cell culture conditions

24 well plates were coated with anti-CD3/28 stimulating antibodies (Biolegend Catalog Nos. 100302 and 102102) at final concentration of 5 µg/mL each overnight at 4° C. The next day, 2 million single cell splenocytes were plated in 1 mL T_RM_-like culture media onto each well with 7.5 ng/mL human IL-2 (Obtained from the NIH). Two days later, cells were collected, centrifuged at 424xg at 4° C for 6 minutes, and plated onto non-coated 24 well plates in 1 mL TRM-like culture media with 7.5 ng/mL IL-2 for three days. After that, cells were collected, centrifuged at 424xg at 4° C for 6 minutes, and plated in TRM-like culture media with 10 ng/mL mouse IL-7 (Biolegend Catalog No. 577804) and 10 ng/mL human TGFβ1 (Fisher Scientific PHG9214) for 6 days.

Additional glucose (Fisher Scientific Catalog No. A2494001) and lipid supplement (Millipore Sigma Catalog No. L5146) were added to TRM-like cell culture media starting from the day of spleen harvest (day 0) throughout the end of the experiment (day 11, day 6 post TGFβ1 addition). Soluble lipid stores were created in Kolliphor P 188 (Millipore Sigma K4894) according to lipid supplement instructions.

### 2.14 Staining for Flow Cytometry

Cells were suspended in 100 µL staining buffer for 30 minutes at 4° C. Staining buffer was created by adding staining antibodies to PBS below for a final concentration of 1 µg/mL each, or 0.2 µL of live-dead stock solution/100 µL staining buffer. Live-dead stock solution was prepared according to package insert available on the vendor website. αCD16/32 was added for a final concentration of 2.5 µg/mL. αCD45 was previously injected at time of organ harvest intravenously at a concentration of 0.2 µg stock antibody/100 µL PBS in a volume of 100 µL. See table S1 for more information. All antibodies used Bangs Laboratories compensation beads (Bangs #556) which were stained an equivalent time. Live-dead stain used Life Technologies ArC reactive beads (Invitrogen A10346). Because the vast majority of cells at day 14 post infection will have been antigen experienced, CD44 was not used as a marker on day 14 experiments. Experiments were run on an 18-color BD LSR II flow cytometer. Gating strategies are displayed in in the supplementary file.

### 2.15 MFI Determination and Normalization

The four subsets of interest were manually gated after selecting for lymphocytes (FSC-A and SSC-A), singlets (FSC-A and FSC-H), live cells, IV-CD45^neg^ cells, TCRβ^high^and CD8α^high^ cells. DN (Double Negative) was defined as CD103^low^ and CD49a^low^, DP (Double Positive) as CD103^high^ and CD49a^high^, CD103 SP (CD103 Single Positive) as CD103^high^ CD49a^low^, and CD49a SP (CD49a Single Positive) as CD49a^high^ and CD103^low^.

Median fluorescence intensities (MFIs) were calculated using Flowjo (version 10.6.2). Data was normalized by dividing the MFI of each subset by the sum of all four subsets’ MFIs for each mouse and multiplying by 4. For the assays using TMRE, FCCP was subtracted from the TMRE MFI for each subset within each mouse. That number was divided by the sum of MFIs of the uncoupled control (FCCP+TMRE) subsets for each mouse. Normalized data can be found at https://github.com/sportiellomike.

### 2.16 Statistical Tests

After repeated measures ANOVA, Multiple pairwise comparisons were evaluated using the Student’s T-test with Benjamini-Hochberg correction. If P_adjusted_<0.05, results were deemed significant. R Packages ggplot2 (version 3.3.2) and rstatix (0.7.0) were used for data plotting and statistical testing respectively. The code used to perform statistical tests and generate figures can be found here: https://github.com/sportiellomike.

### 2.17 Ethics statement

This study was carried out in strict accordance with the recommendations in the Guide for the Care and Use of Laboratory Animals as defined by the NIH. Animal protocols were reviewed and approved by the Institutional Animal Care and Use Committee of the University of Rochester in writing. All animals were housed in a centralized and Association for Assessment and Accreditation of Laboratory Animal Care accredited research animal facility that is fully staffed with trained husbandry, technical, and veterinary personnel. Our UCAR number is 2006-029. Euthanasia was performed by animal welfare committee reviewed approved methods.

## 3 Results

### 3.1 Pathway analysis for metabolic analyses

T_RM_ can be defined by their surface protein expression, with multiple markers having been used including CD69, CD49a, and CD103. In the mouse lung, CD44 expression identifies antigen-experienced memory T cells while CD49a marks T_RM_, and when used in combination with CD103, yields two T_RM_ subsets: CD49a^pos^CD103^pos^ (<u>D</u>ouble <u>P</u>ositive, DP) and CD49a^pos^CD103^neg^ (<u>CD49a</u> <u>S</u>ingle <u>P</u>ositive, CD49a SP) (7). At later memory time points, very few CD49a^neg^CD103^pos^ (<u>CD103 S</u>ingle <u>P</u>ositive, CD103 SP) cells remain, suggesting they have either died, left the tissue, or altered their integrin expression. CD49a^neg^CD103^neg^ (<u>D</u>ouble <u>N</u>egative DN) cells represent mixed phenotypes, though more than half are CD44^high^ CD62L^low^ T cells consistent with an Effector Memory profile (T_EM_) and may be circulating (7). To more fully describe the metabolic profiles of T_RM_ based on integrin expression, we utilized the publicly accessible RNA sequencing data set we previously reported (7). This data set was generated from CD44^high^ mouse lung tissue CD8 T cells 21 days post-infection (dpi) that were sorted by CD49a and CD103 expression.

Using *P_adj_* <0.05 as a cutoff, the data were analyzed to identify genes with a minimum of Log_2_(x) = 0.5 Fold Change (LFC) to determine up- or downregulated genes, respectively. Per pathway analysis utilizing both the Reactome and KEGG databases, there were no significant differences when comparing DP to CD49a SP. However, several metabolic pathways were upregulated in DP vs DN, including *Metabolism of lipids and lipoproteins, Fatty acid, triacylglycerol, and ketone body metabolism*, and *Cholesterol biosynthesis.* A selection of pathways is plotted in **Figure 1A-B**. The complete set of significantly enriched pathways can be found in the Supplementary file “Significantly Enriched Pathways.xlsx”.

**Figure 1.**
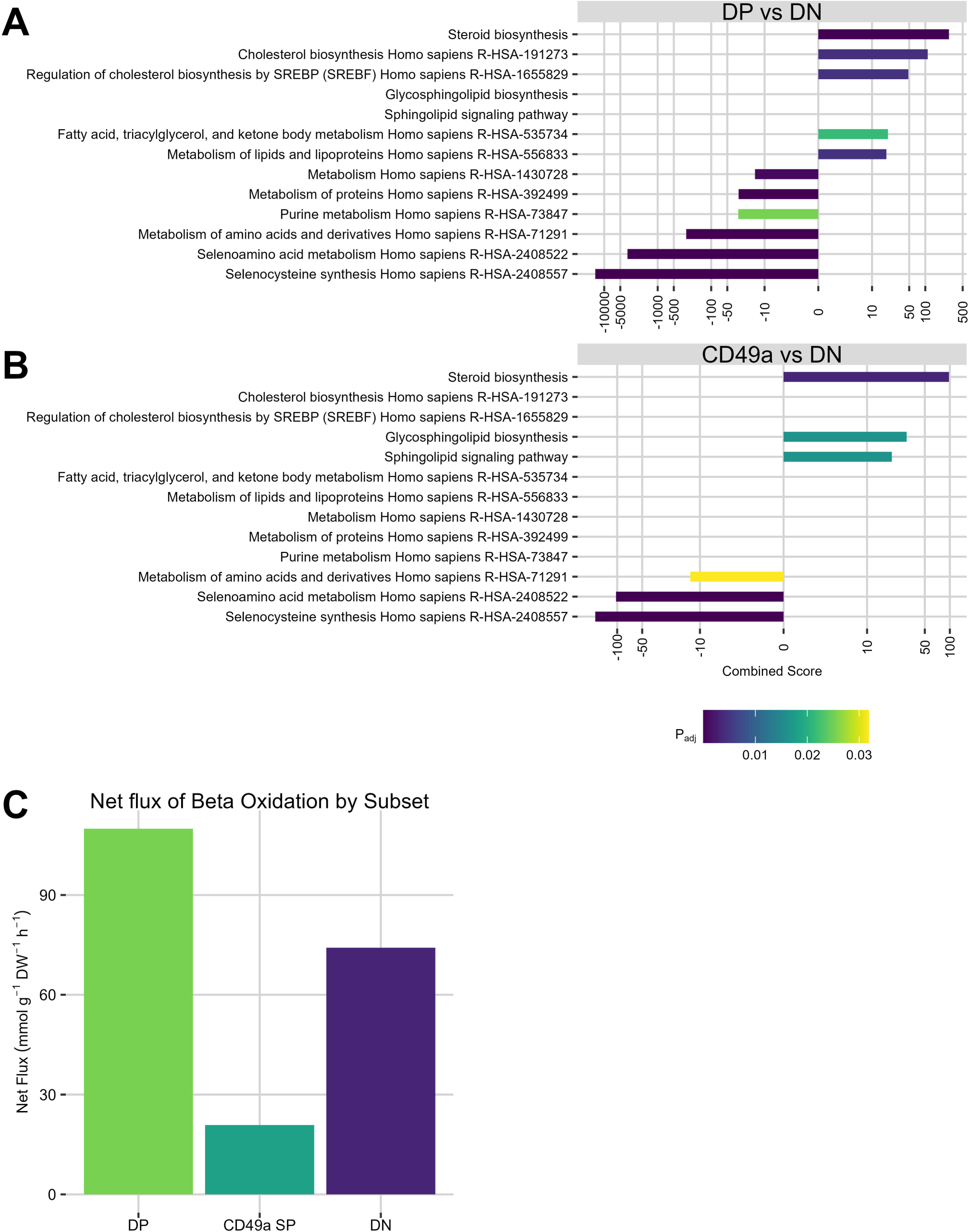
Transcriptomic analysis of lung CD8 T cells 21 days post infection. Reactome and KEGG databases were queried with the lists of differentially expressed genes between (A) DP and DN,or (B) CD49a SP and DN. Criteria for inclusion as differentially expressed genes include P_adj_ < 0.05 and Log_2_(Fold Change) > 0.5 or Log_2_(Fold Change) < -0.5. Adjusted P value cutoff for enrichment is P_adj_<0.05. Only select pathways are shown here. A complete list can be found in the supplemental. Datasets were generated using 10 total mice per N with N=3 per subset. Metabolic models were created using the GIMME algorithm and the level of flux through Beta Oxidation was plotted (C). DP (<u>D</u>ouble <u>P</u>ositive, CD49a^pos^CD103^pos^); DN (<u>D</u>ouble <u>N</u>egative, CD49a^neg^CD103^neg^); CD49a SP (<u>CD49a</u> <u>S</u>ingle <u>P</u>ositive, CD49a^pos^CD103^neg^).

Interestingly, CD49a SP and DP did not share any strictly metabolic pathways from the Reactome when compared to DN yet were indistinguishable from one another at the global transcriptomic level (7) (**Figure 1B**). They also did not share any pathways with each other in the KEGG database beyond *Steroid biosynthesis* (**Figure 1B**). Metabolic differences between DP and DN but not CD49a SP and DN could imply metabolic differences between CD49a SP and DP that these pathway analysis methods are not sensitive enough to distinguish. In summary, T_RM_ subsets defined by integrin expression were metabolically distinguishable from DN cells based on gene expression.

### 3.2 Metabolic modeling predicts lipid-centric metabolism in T_RM_

Having performed basic pathway analyses and discovering a possible enrichment for lipid-centric programming in T_RM_, we developed an approach for modeling the metabolome of lung CD8 T cells using the entirety of the transcriptomic data and used this more intensive approach to verify the results of the basic analysis.

Metabolic models are networks of metabolites and the reactions that utilize and produce them. Generally, these reactions are catalyzed using genome-encoded enzymes. Three metabolic computational models were constructed: one for lung CD8 T cells expressing both CD49a and CD103 (*tissuemodelDP*), one for the single positive subset (*tissuemodelCD49aSP*), and one for the subset expressing neither integrin (*tissuemodelDN*). With these models, we used the expression of enzymes present in a system to predict the amount of each metabolite present. These models together incorporate nearly all antigen-experienced CD8 T cells in mouse lungs, encompassing both T_RM_ and non-T_RM_ memory T cells.

Briefly, the published mouse metabolic model iMM1865 was used as a starting point (12). Then, the Gene Inactivity Moderated by Metabolism and Expression (GIMME) algorithm was implemented in the COBRA Toolbox (13) to conduct a flux balance analysis using our RNAseq transcript level data to inform the level of flux for each reaction (10). GIMME alters the level of flux through a reaction to minimize the total inconsistency score. Assuming a larger quantity of enzyme should produce a larger flux through that reaction, GIMME increases (or decreases) flux through different reactions depending on the level of transcript for that enzyme. When building metabolic models, an objective function must be selected for which flux is to be maximized given the constraints. In line with most other metabolic models, “BIOMASS” was selected, which itself is a combination of several reactions related to cell maintenance that must have positive flux for cells to be viable. Given our constraints, a single, nonzero optimal solution was predicted for all three of our models.

Our model predicted that our DP subset had the greatest flux through its *Beta Oxidation of Fatty Acids* pathway, consistent with our initial pathway analysis (**Figure 1C**). In addition, the CD49a SP subset differed from the DP subset with less flux through that pathway than both DP and DN, again consistent with the initial pathway analysis. These results, in addition to our basic pathway analysis, imply differences in T_RM_ metabolism between DP and CD49a SP subsets, as well as between integrin-expressing and DN subsets.

### 3.3 T_RM_ CD8 T cell subsets have unique metabolic profiles directly ex vivo

Thus far, our transcriptomic analyses were largely consistent with that demonstrated for bulk T_RM_ in the skin. Nonetheless, transcriptomic data and metabolic modeling do not clearly demonstrate that functional differences exist between cell subsets whether circulating or tissue memory. The Seahorse XF Mito Fuel Flex assay which (ideally) requires large cell numbers is considered a “gold standard” for measuring a number of metabolic metrics such as oxygen consumption and acidification rate. However, due to limiting cell numbers and our interest in the resting (not restimulated) metabolic state of memory cells, the signal was not high and reproducible enough to draw conclusions, demonstrating this approach is not feasible with unstimulated primary memory T cells. Instead, to capture single-cell data, a flow-cytometric approach was performed. T cells with T_RM_-like integrin expression are evident in mouse lung T cells 14 days post-infection (dpi) with HKx31 influenza A virus and may include cells with memory potential, while by 60 dpi memory is well established. To determine if the T cell subsets display any differences in their mitochondria, the mitochondrial mass of cells of different integrin profiles was evaluated by Mitotracker Green FM (Mitotracker) staining. Mitotracker preferentially accumulates in mitochondria by covalently binding free thiol groups independent of mitochondrial membrane potential (14). Both airway cells harvested through bronchoalveolar lavage (BAL) and cells remaining in the tissue after lavage (likely lung parenchymal cells) were examined for Mitotracker staining. Tissue vasculature-associated cells were excluded using a standard intravenous (IV) labeling approach (7).

We found no differences in Mitotracker staining in the lung or BAL at day 14 (**Figure 2A and S2**). However, at day 60 dpi, CD49a SP had significantly lower levels of mitochondrial mass than DP and DN counterparts in the lung, while DP had the highest staining (**Figure 2B**). This suggests that as memory T cell subsets develop, mitochondrial mass changes.

**Figure 2.**
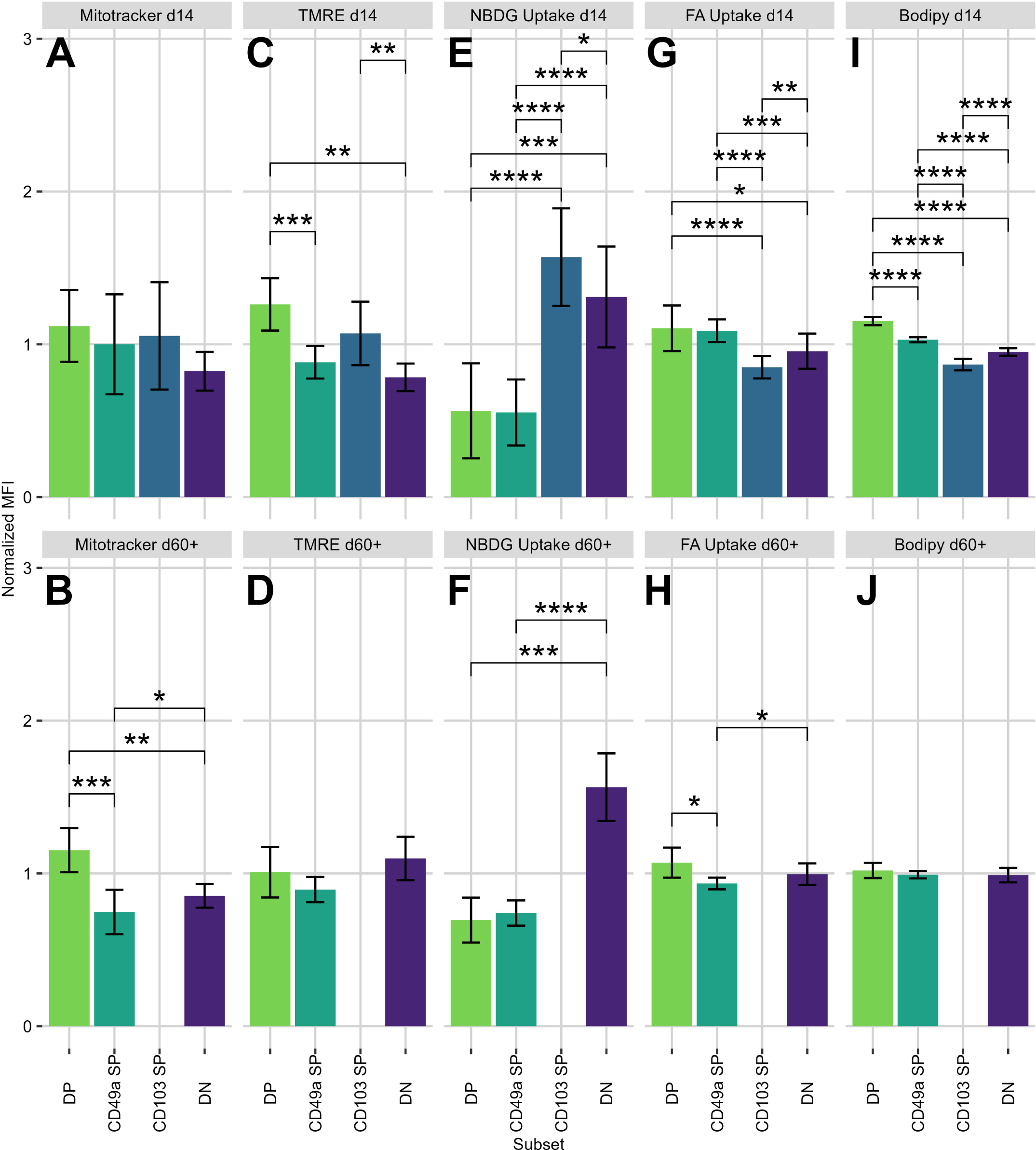
Normalized mouse IV-label-negative lung CD8 T cell MFIs from flow cytometric assays. Normalized MFIs of IV-label-negative lung CD8 T cells plotted for each assay at 14 and 60+ days post infection derived from flow cytometric analysis. *P < 0.05, **P < 0.01, ***P < 0.001, ****P < 0.0001. DP (<u>D</u>ouble <u>P</u>ositive, CD49a^pos^CD103^pos^); DN (<u>D</u>ouble <u>N</u>egative, CD49a^neg^CD103^neg^); CD49a SP (<u>CD49a</u> <u>S</u>ingle <u>P</u>ositive, CD49a^pos^CD103^neg^) ; CD103 SP (<u>CD103</u> <u>S</u>ingle <u>P</u>ositive, CD49a^neg^CD103^pos^). Mitochondrial mass was assessed with Mitotracker Green FM (A and B). Mitochondrial membrane potential was assessed with TMRE (C and D). Glucose uptake was assessed with glucose analogue 2-NBDG (E and F). Fatty acid uptake was assessed with fluorescently conjugated fatty acids (G and H). Neutral lipid stores were assessed with Bodipy (I and J). Data generated from two independent experiments of N= 4-10 mice per experiment.

The question remained of whether the cell populations displayed differential mitochondrial functions, not just different mitochondrial masses. Oxidative phosphorylation is a primary role of the mitochondrion, as it is the most efficient means by which a cell can generate ATP. Cells with equivalent mitochondrial mass could potentially undergo differential rates of oxidative phosphorylation and therefore ATP production. Oxidative phosphorylation is linked to mitochondrial membrane polarization. TMRE, a fluorescent dye that preferentially stains polarized mitochondrial membranes was used as a proxy for processes that polarize the mitochondrion, though TMRE is not a direct readout of oxidative phosphorylation. To remove background staining, an equivalent sample was treated with both TMRE and the decoupling agent FCCP, whose signal was subtracted from the treatment group’s signal.

Cells in the lung expressing CD103 had increased TMRE staining implying a greater degree of membrane polarization and therefore oxidative phosphorylation 14 dpi (**Figure 2C**). Later, at 60+ dpi, no subset was significantly different (**Figure 2D**). At 60+ dpi, DP and DN cells have more mitochondrial mass than CD49a SP cells (**Figure 2B**), while having the same membrane potential (**Figure 2D**). Though not the only possibility, this is consistent with cells that have a higher spare respiratory capacity (SRC), i.e., they may have the capability to use their higher amounts of mitochondria to produce more ATP after reactivation.

Based on the observed differences, we wanted to investigate whether nutrient uptake was altered in a similar way. Glucose is a metabolically essential molecule in a variety of cellular processes as both a source of cell energy in the form of ATP as well as a carbon source in anabolism. To evaluate the extent to which glucose was being taken up by the CD8 T cells, we utilized the glucose analog 2-NBDG. This fluorescent analog undergoes facilitated diffusion through glucose transporters GLUT1-4 as well as through active transport through SGLT1 and 2, recapitulating the kinetics of glucose transport (15, 16, 17, 18, 19, 20). It is phosphorylated by hexokinase after entering the cell, which traps the fluorescent molecule in the cell in a form that cannot proceed further in glycolysis.

Lung DP and CD49a SP displayed similar levels of 2-NBDG staining 14 dpi, however, these populations displayed lower levels of staining compared with DN and CD103 SP cells, suggesting that CD49a expressing subsets may be less glucose-dependent than CD49a negative (non-T_RM_) counterparts (**Figure 2E**). A similar pattern was observed in BAL samples, with CD49a^pos^ subsets taking up lower levels of the dye (**Figure S2**). This pattern persisted at later memory time points 60+ dpi (**Figure 2F**) and is consistent with studies done on T_RM_ in the skin, which noted an increased dependence on free fatty acids (FFAs) as opposed to glucose (9).

Having demonstrated significant differences in the uptake of the glucose analog 2-NBDG, we next investigated the ability of T cells to take up free fatty acids with the hypothesis that lung T_RM_ utilize a more lipid-centric metabolism, similar to skin (9). A fluorescently conjugated fatty acid (FFA) was utilized to measure uptake. In the lung parenchyma, CD49a^pos^ subsets (CD49a SP and DP) took up the highest amount of FFA. (**Figure 2G**). Similar results were found among the BAL subsets. These findings were similar but distinct at 60+ dpi: DP and DN took up more FFAs than CD49a SP (**Figure 2H**). Other studies that have reported a dependence of T_RM_ on FFAs have not used CD49a as a marker (9) and therefore may have excluded the CD49a^pos^ CD103^neg^ subset while possibly including the CD103 SP subset. We expected our results to demonstrate that both DP and CD49a SP (the latter not studied by other metabolic studies) were equally dependent on these FFAs in the lung, but we found this only to be true at 14 dpi. At 60+ dpi, DP were much more like DN than their CD49a SP T_RM_ counterparts. One interpretation consistent with the data is that CD49a SP utilizes a less lipid-centric metabolic strategy than their DP counterparts.

We next wanted to determine if there were further differences in metabolic lipids. To address this, Bodipy, a molecule that is taken up passively and stains cellular stores of neutral lipids, was used to quantify the amount of neutral lipid present in lung T cells (21). Neutral lipids serve as a store of potential energy as well as a carbon source, a valuable resource if the cells were to expand and undergo effector functions in a nutrient-deficient environment.

Nearly identical patterns for BAL and lung were observed at 14 dpi (**Figure 2I, Figure S2**). DP cells displayed the highest levels of neutral lipid storage, followed by CD49a SP then DN, with the lowest levels in CD103 SP. While all six comparisons demonstrate that the four subsets are significantly different from one another, it is remarkable that the subsets expressing CD49a (CD49a SP and DP) stain brightly for neutral lipids while those not expressing CD49a (CD103 SP and DN) display low levels. Having quick access to neutral lipids for both energy and carbon needs may prove important for their protective function and/or long-term memory formation given these differences observed at 14 dpi were not seen at 60+ dpi (**Figure 2J**).

### 3.4 Glucose and lipid metabolites alter the phenotype of T_RM_-like cells in vitro

Noting the myriad of differences between T_RM_ and non-T_RM_, as well as among T_RM_ subsets, we hypothesized that the metabolic microenvironment is deterministic in their differentiation. Given that tissue-specific metabolic substrates are impossible to control, an *in vitro* approach was employed. T_RM_ differentiation in vivo requires TGFβ (22). Similarly, T cells with T_RM_-like integrin profiles can be generated by stimulating naïve mouse splenocytes under defined conditions that include TGFβ (23). Briefly, T cells were initially activated by culture on αCD3/CD28 coated plates for two days in the presence of IL-2, followed by three days in non-coated plates supplemented with IL-2. T_RM_-like differentiation was achieved by placing these cells in IL-7 and TGFβ for six more days to elicit a T_RM_-like surface phenotype (23). To test the effects of glucose or lipids within as well as across levels, both the concentration of glucose as well as lipids were independently varied (**Figure S3A**). Glucose concentrations began with the concentration of glucose in standard cell culture media (4.5 mg/ml). This concentration has been demonstrated to ensure optimal T cell viability and activation, especially over the course of several days in culture since glucose is rapidly utilized by cells, though it is higher than that found in the blood or most other tissue microenvironments. To assess the impact of lipids on T-cell differentiation *in vitro*, a broad range of exogenous lipid concentrations, which includes both fatty acids and cholesterol (Millipore Sigma #L5146), was used. Lipid supplementation varied from no exogenous lipid to 8x the concentration often used for cell culture.

First, the effect on CD49a expression was examined. Without exogenous lipid added, there was no effect of glucose concentration on frequency, but with 2x or 4x lipid supplementation, generally, as the concentration of glucose increased, the cell number (normalized per volume) and proportion of CD49a positive cells both decreased (**Figures 3A-3B, S3, Supplementary File “2D combined stats sheet.xlsx”**). The relationship between the absolute number of CD49a positive cells per volume and glucose concentration was linear. 8x lipid supplementation was toxic as the number of live CD8 T cells for the group was effectively zero. Overall, this paints a picture of increasing glucose concentration causing a decrease in both the frequency and the absolute number of CD49a-positive cells. This contrasts with the frequency of central memory marker CCR7 positivity (**Figures 4A, 4B, and S3, Supplementary File “2D combined stats sheet.xlsx”**), which did not vary with lipid supplementation, though increasing concentrations of glucose did seem to decrease the frequency of CCR7 positive cells.

**Figure 3.**
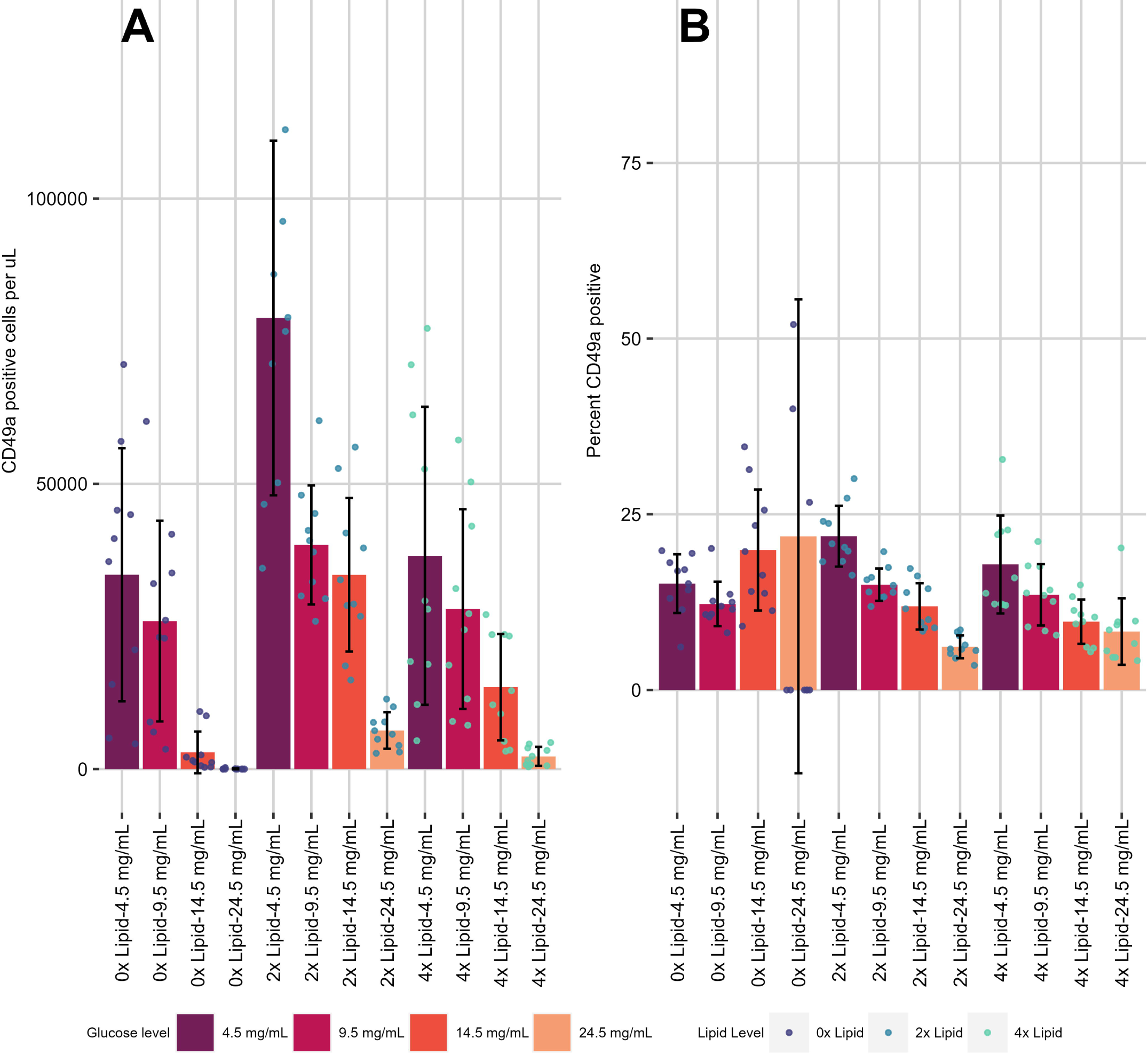
Metabolite concentration impact on T_RM_-like cell differentation and maintenance *in vitro*. CD49a positive cells per µL (A) and percent CD49a positive (B) were assessed in metabolite alteration assays where both lipid (0x-4x) and glucose (4.5-24.5 mg/mL) were altered after activation in the presence of IL-2 and cultured with IL-7 and TGFβ. *P < 0.05, **P < 0.01, ***P < 0.001, ****P < 0.0001. Small circles represent individual data points. Data generated from two independent experiments of N=5 mice per experiment.

**Figure 4.**
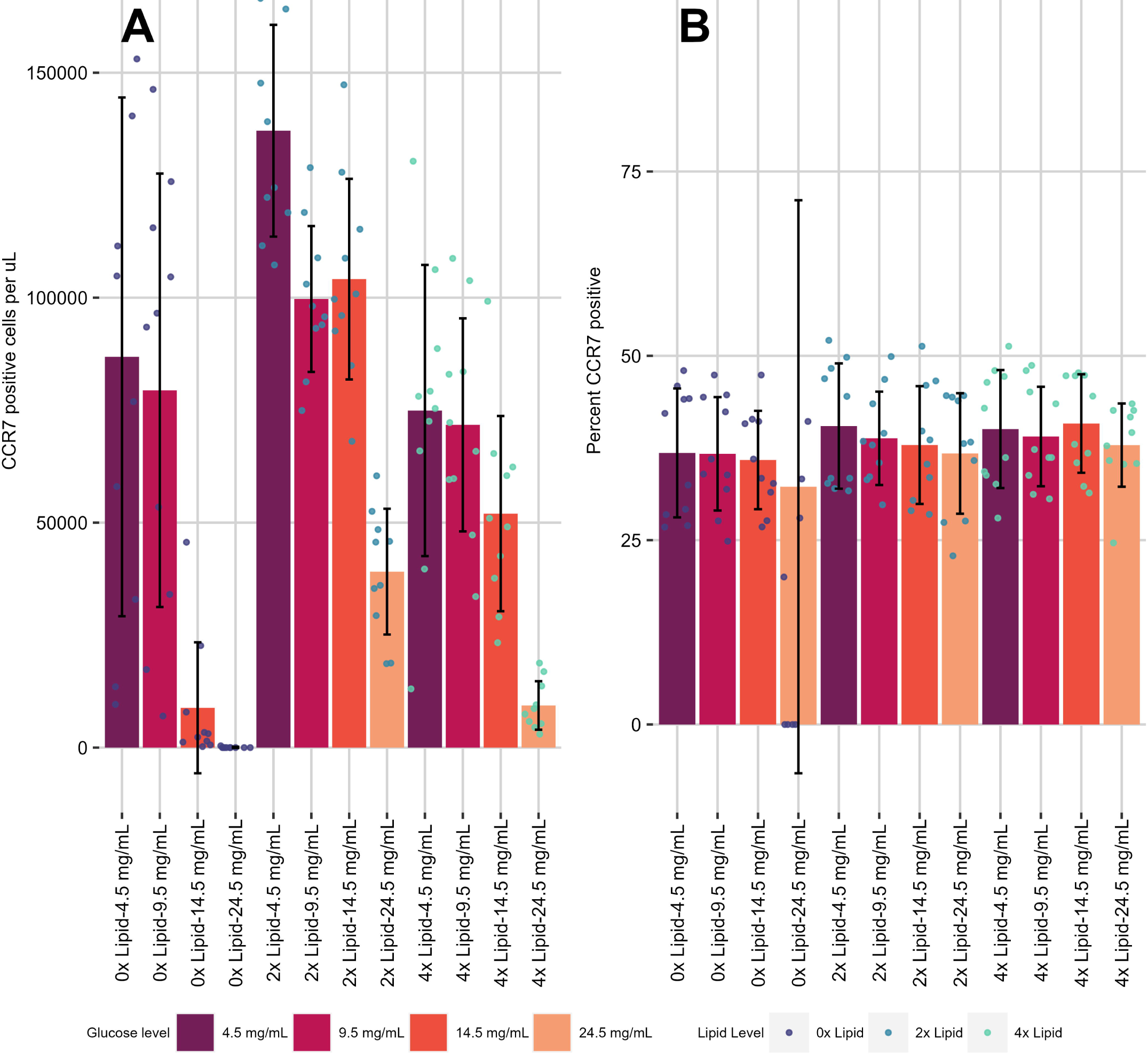
Metabolite concentration impact on T_CM_-like cell differentation and maintenance *in vitro*. CCR7 positive cells per µL (A) and percent CD49a positive (B) were assessed in metabolite alteration assays where both lipid (0x-8x) and glucose (4.5-24.5 mg/mL) were altered after activation in the presence of IL-2 and cultured with IL-7 and TGFβ. *P < 0.05, **P < 0.01, ***P < 0.001, ****P < 0.0001. Small circles represent individual data points. Data generated from two independent experiments of N=5 mice per experiment.

Looking at the effect of lipid supplementation at constant glucose levels on the differentiation of CD8 T cells *in vitro*, a contrasting pattern emerged: within each concentration of glucose a 2x lipid supplement increased the frequency of CD49a positive cells at the lower concentrations of glucose tested (4.5 and 9.5 mg/mL) (**Figure 3B, Supplementary File “2D combined stats sheet.xlsx”**). Further addition of lipid to 4x did not increase the frequency of CD49a positive cells. The absolute number of CD49a positive cells per volume peaked at 2x and decreased upon further addition of lipid for all concentrations of glucose tested. A similar but less drastic finding occurred for CCR7 positive cells, but not for the frequency of positive cells (**Figure 4B, Supplementary File “2D combined stats sheet.xlsx”**). Patterns for CD49a^pos^CD103^pos^ cells were similar to those of CD49a SP (**Figure S5**).

Taken together, these results imply that increasing glucose availability antagonizes the generation or maintenance of memory CD8 T cells independent of lipid supplementation and affects the expression of CD49a, at least in this assay. Furthermore, lipid supplementation is beneficial for CD49a expression up to a point but can be toxic at too high a level.

## 4 Discussion

Using a genomic approach, we found memory T cell subsets in the lung exhibit distinct metabolic profiles. Furthermore, *in vitro* experiments conducted to drive T cells towards T_RM_-like surface phenotypes in the presence of varying metabolic substrates suggest metabolite availability can influence memory T cell differentiation and/or survival. *In vivo* metabolite availability may be due to differences in local concentrations, the cell’s ability to transport metabolic substrates into the cell, or the expression of receptor(s) needed to take up these metabolites from outside the cell.

Differential expression, pathway, and metabolic modeling of our RNAseq data led to the hypothesis that lung T_RM_ cells have distinct metabolic programming compared with circulating cells. Using both functional and descriptive methods, we produced a dataset for a variety of cell functions including mitochondrial profiling and nutrient uptake at both 14 and 60+ dpi. Integrating these results, what seems clear is that DP and CD49a SP, though different from each other in a few assays, share a similar pattern. This pattern is distinct from DN T cells, a mixed population consisting mostly of T cells with the phenotypes of effector memory and some central memory (7). The methods employed suggest DP and CD49a SP both take up more fatty acids, store more neutral lipids, and take up less glucose than DN cells at 14 dpi. At later memory time points, DP and CD49a SP diverge: DP cells have more mitochondrial mass and take up less glucose and more fatty acid than CD49a SP. Generally, memory cells have a higher SRC than effector cells, suggesting one possible advantage is to convey spare respiratory capacity (24). An increased spare respiratory capacity in DP but not CD49a SP could imply different roles in memory and effector responses. This should be cautiously interpreted since this SRC was not directly measured here, meriting further studies to validate this conclusion.

These findings are consistent with the interpretation that DP cells exhibit a more typical “resting” memory phenotype and that CD49a SP could represent a more “activated” cell type. We have previously reported that the CD49a SP subset has a number of more effector-like features such as having a higher frequency of Granzyme B and perforin double positive cells, making them poised for cell killing (7).

The metabolic profile of an individual T cell as well as a T cell subset is important functionally in differentiation as well as effector function. Indeed, this is a leading hypothesis as to why some T cells become T_RM_ and some do not (23). The functions of these subsets during a subsequent infection may in part be achieved through the availability of different nutrients: if T_RM_ depend mostly on the uptake of free fatty acids whereas naive and effector T cells depend more on glucose uptake, both may be able to perform their functions without competing for energy resources. Furthermore, these profiles can directly impact the cell signaling the cells are receiving. It is known that altering the ability of naïve CD8 T cells to take up glucose directly impacts their likelihood of becoming memory cells such that the most glucose-starved T cells become memory cells (25). Having a metabolic profile that, at baseline, is not dependent on glucose may ensure that the internal cell signaling conferring this memory phenotype continues to do so in a positive feedback manner. TCR stimulation is directly related to the specificity of binding and the abundance of a TCR’s peptide:MHC complex. Others have shown that T cells take up more glucose when more activated (26). Having an abundance of high-affinity interactions with peptide:MHC complexes may cause a hyper-local effect of glucose starvation. The same group demonstrated that decreased availability of glucose led to increased memory CD8 T cell populations. A lipid-centric metabolism by memory T-cells may assist naïve T cells to respond to infection because they don’t compete with other memory cells for glucose making it less likely that memory T cells will starve out the naïve responses. This may support eliciting a more diverse group of memory T cells at the end of an infection, which would increase the likelihood of a productive immune response upon subsequent heterologous infection. An important caveat to our interpretation is that NBDG doesn’t always correlate with glucose uptake, even though it is a glucose analog itself. As a result, more research must be done to confirm these conclusions.

The causal effect of metabolic factors on CD8 T cell differentiation has been studied but never in the context of T_RM_ differentiation or in consideration of different T_RM_ subsets. In short, we found glucose to be detrimental to the *in vitro* driven expression of T_RM_ marker CD49a in a linear manner, whereas we found increasing lipid beneficial, to a point. Conversely, glucose did not affect the expression of central memory marker CCR7, whereas the effect of lipids was similar for both CD49a and CCR7. Optimal concentrations may exist that increase or decrease the expression of these markers. We believe these findings have direct implications for the creation of therapies like CAR-T cells which necessitate *ex vivo* culture conditions. For example, conditions could be optimized for the creation of T-cells with a T_RM_-like phenotype as observed among tumor-infiltrating lymphocytes associated with positive outcomes (27, 28, 29, 30). Future studies should assess this possibility.

## Supporting information

Signficantly enriched pathways for Figure 1

Statistics for Figures 3 and 4

## 5 Figure Legends

**Supplementary Figure 1:** Gating scheme and representative fluorescent dye staining for flow cytometry assays pictured in Figure 2.

**Supplementary Figure 2:** Normalized mouse bronchoalveolar lavage non-intravenously labelled CD44^pos^ CD8 T cell MFIs from flow cytometric assays. *P < 0.05, **P < 0.01, ***P < 0.001, ****P < 0.0001. DP (<u>D</u>ouble <u>P</u>ositive, CD49a^pos^CD103^pos^); CD49aSP (<u>CD49a</u> <u>S</u>ingle <u>P</u>ositive, CD49a^pos^CD103^neg^); CD103SP (<u>CD103</u> <u>S</u>ingle <u>P</u>ositive, CD49a^neg^CD103^pos^); DN (<u>D</u>ouble <u>N</u>egative, CD49a^neg^CD103^neg^). Data were generated from two independent experiments of 4-10 mice each.

**Supplementary Figure 3. Assay layout for T cell differentiation studies:** 16 combinations (4 glucose treatment groups, 4 lipid treatment groups permuted) were assayed. Glucose groups included 4.5, 9.5, 14.5, and 24.5 mg/mL. Lipid groups included 0x, 2x, 4x, and 8x recommended concentrations (A). Gating scheme for differentiation studies (B).

**Supplementary Figure 4. Metabolite concentration impact on T cell viability *in vitro*:** Percent alive in metabolite alteration assays where both lipid and glucose were altered. Data were generated from two independent experiments of 5 mice each.

**Supplementary Figure 5. Metabolite concentration impact on CD49a^pos^CD103^pos^ T_RM_-like cell differentiation and maintenance *in vitro*:** Percent CD49a^pos^CD103^pos^ (B) and cell number per µL (A) in metabolite alteration assays where both lipid (0x-4x) and glucose (4.5-24.5 mg/mL) were altered after activation in the presence of IL-2 and culture with IL-7 and TGFβ. *P < 0.05, **P < 0.01, ***P < 0.001, ****P < 0.0001. Dots represent individual mice. Data were generated from two independent experiments of 5 mice each.

**Supplementary Table 1: Antibody information and catalog numbers.** Additional information to ensure reproducibility in methods.

**Significantly Enriched Pathways.xlsx:** The complete set of significantly enriched pathways from which a subset are plotted in Figure 1 is contained in this table.

**2D combined stats sheet.xlsx:** The statistical comparisons for figures 3, 4, and Supplementary figure 5 are contained in this table.

## 6 Author Contributions

MS designed and performed experiments, analyzed and interpreted data, and wrote the manuscript. AP assisted in conducting experiments and assay optimization. AG, KL, ECR, and TJ edited the manuscript. ECR and KL provided technical support. DJT provided overall direction, procured funding, interpreted data, and edited the manuscript.

## Acknowledgments

We acknowledge the important contributions of the University of Rochester Medical Center Genomic Research Center, the Flow Cytometry Core, and Vivarium. In addition, we are gracious for the reagents provided by Drs. Andrea Amitrano and Minsoo Kim.

## 7 Data Availability Statement

The datasets for this study can be found on GitHub (https://github.com/sportiellomike/Lung-CD8-T-cell-Immunometabolism). Previously published RNAseq data sets were used and are available here: Geo with accession GSE179653 (https://www.ncbi.nlm.nih.gov/geo/query/acc.cgi?acc=GSE179653).

