## Supplementary Figures and Supplementary Table 1 for "Mouse Memory CD8 T cell subsets defined by Tissue Resident Memory (T_RM_) Integrin Expression Exhibit Distinct Metabolic Profiles"

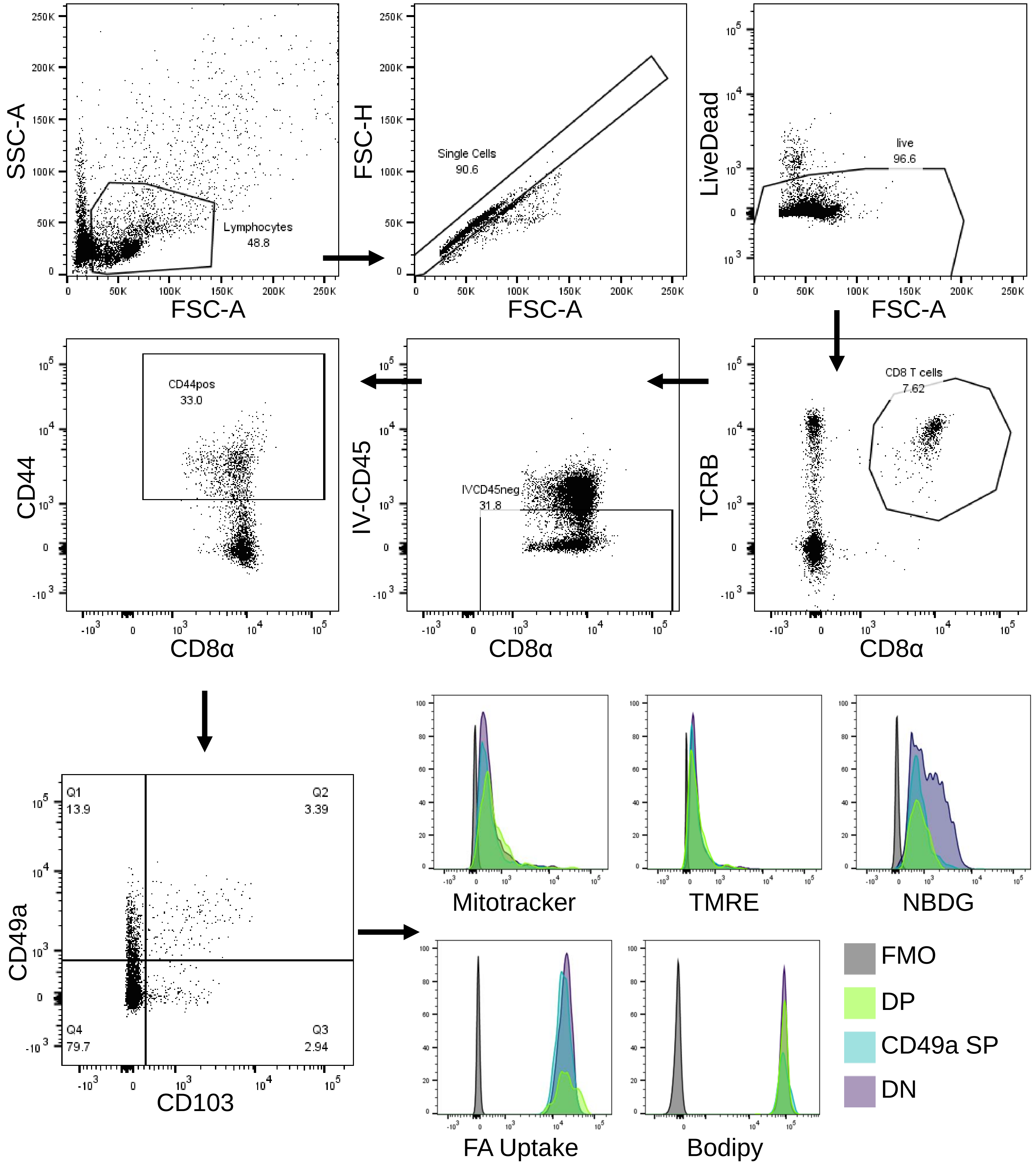


**Supplementary Figure 1:** Gating scheme and representative fluorescent dye staining for flow cytometry assays pictured in Figure 2.

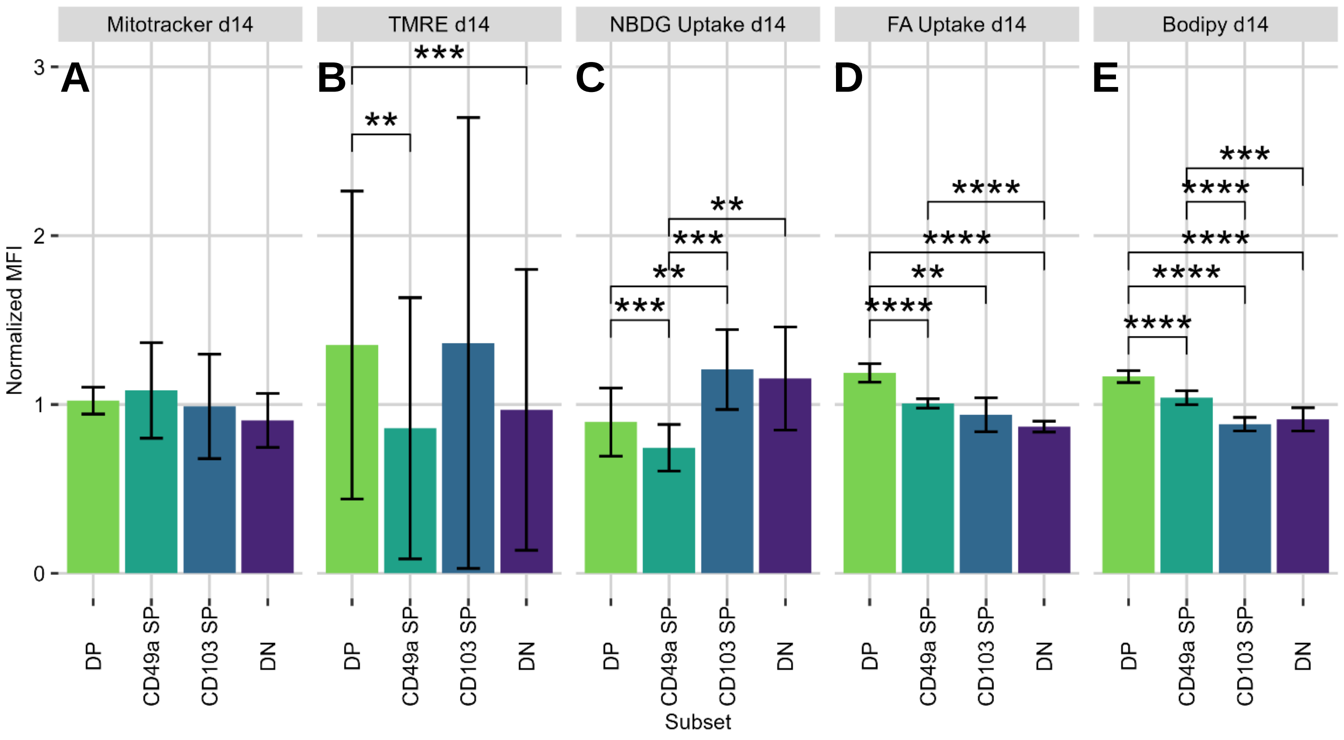


**Supplementary Figure 2:** Normalized mouse bronchoalveolar lavage non-intravenously labelled CD44$^{pos}$ CD8 T cell MFIs from flow cytometric assays. *P < 0.05, **P < 0.01, ***P < 0.001, ****P < 0.0001. DP (Double Positive, CD49a$^{pos}$CD103$^{pos}$); CD49aSP (CD49a Single Positive, CD49a$^{pos}$CD103$^{neg}$); CD103SP (CD103 Single Positive, CD49a$^{neg}$CD103$^{pos}$); DN (Double Negative, CD49a$^{neg}$CD103$^{neg}$). Data were generated from two independent experiments of 4-10 mice each.

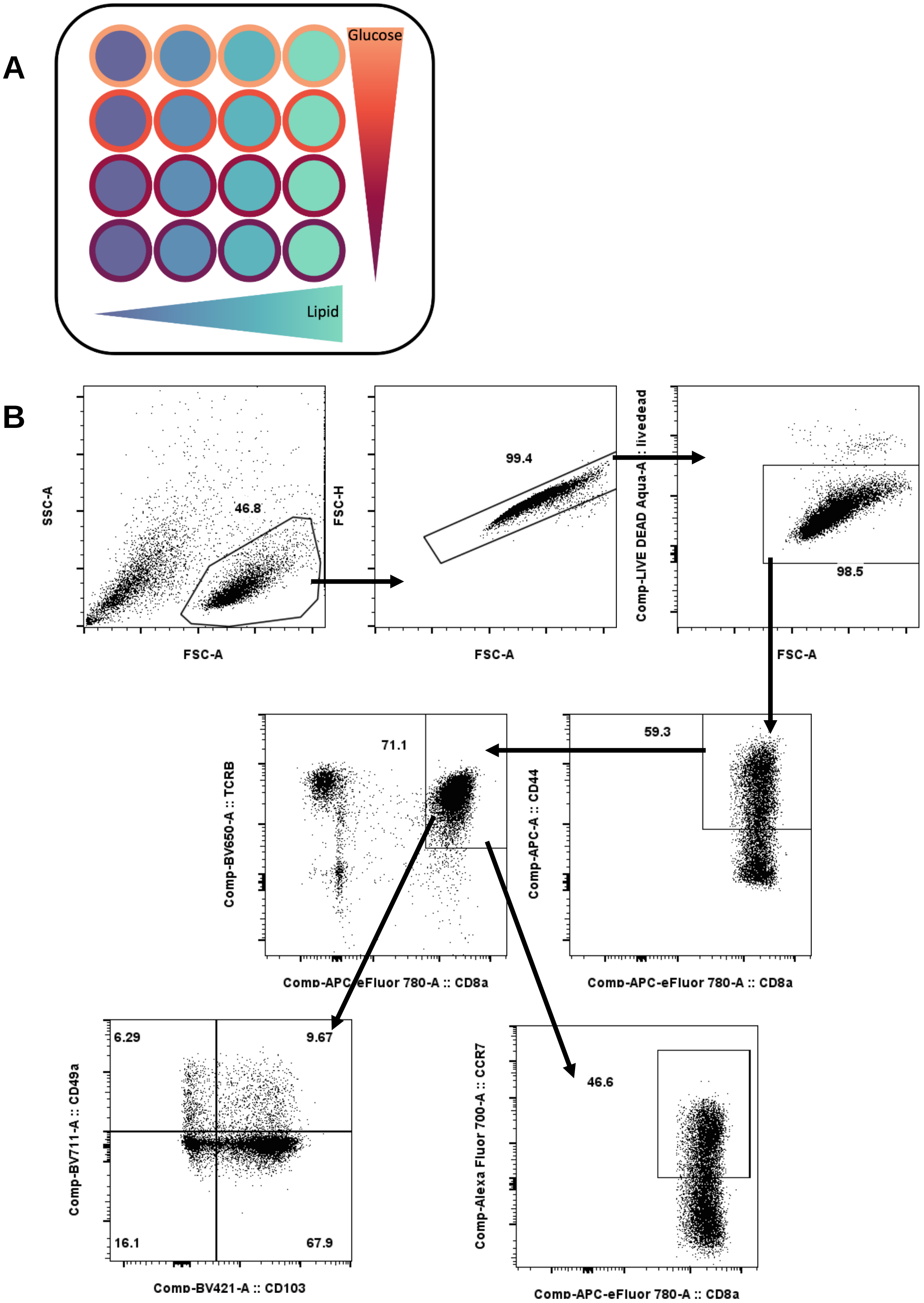


**Supplementary Figure 3. Assay layout for T cell differentiation studies:** 16 combinations (4 glucose treatment groups, 4 lipid treatment groups permuted) were assayed. Glucose groups included 4.5, 9.5, 14.5, and 24.5 mg/mL. Lipid groups included 0x, 2x, 4x, and 8x recommended concentrations (A). Gating scheme for differentiation studies (B).

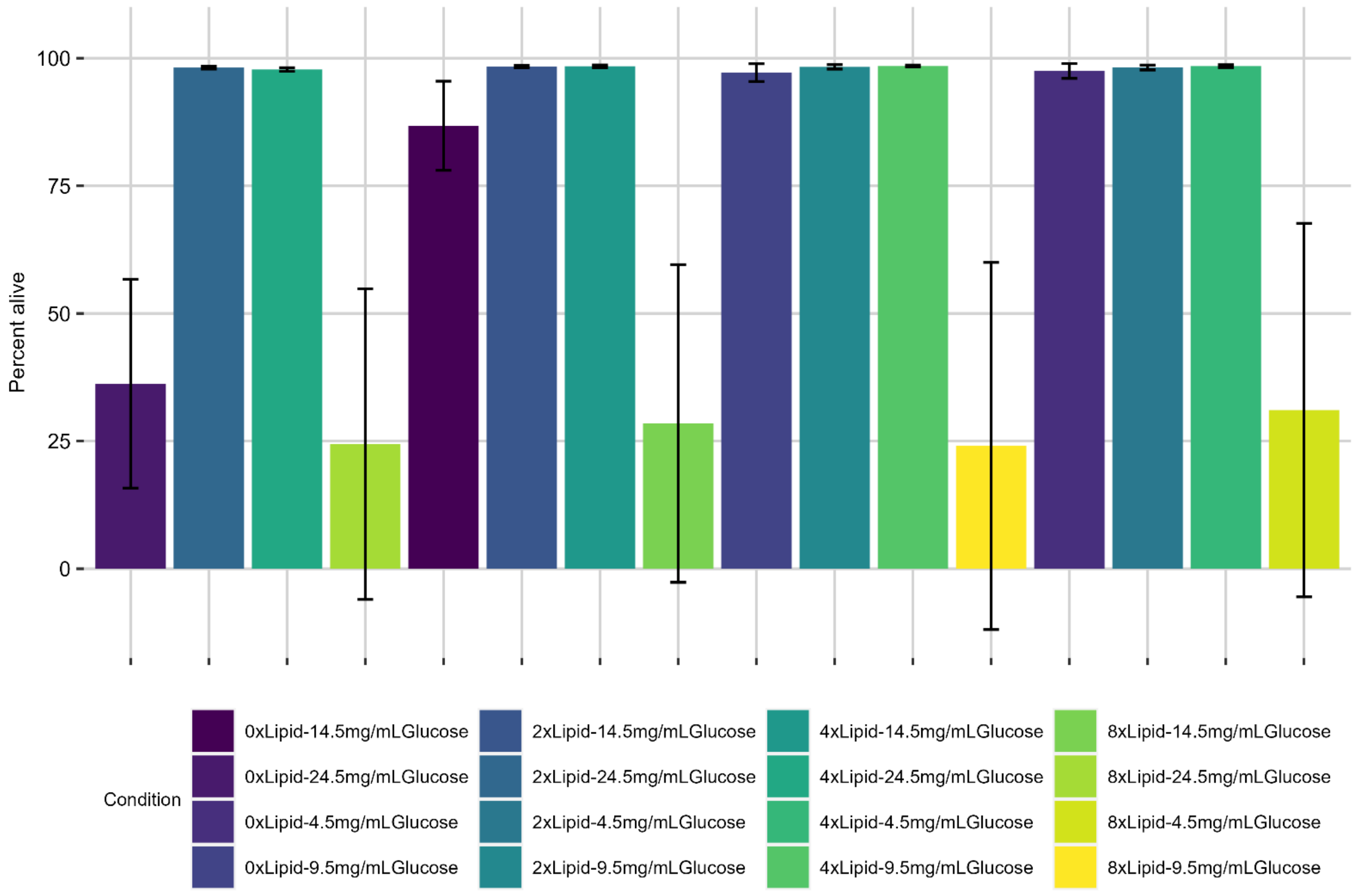


**Supplementary Figure 4. Metabolite concentration impact on T cell viability *in vitro*.** Percent alive in metabolite alteration assays where both lipid and glucose were altered. Data were generated from two independent experiments of 5 mice each.

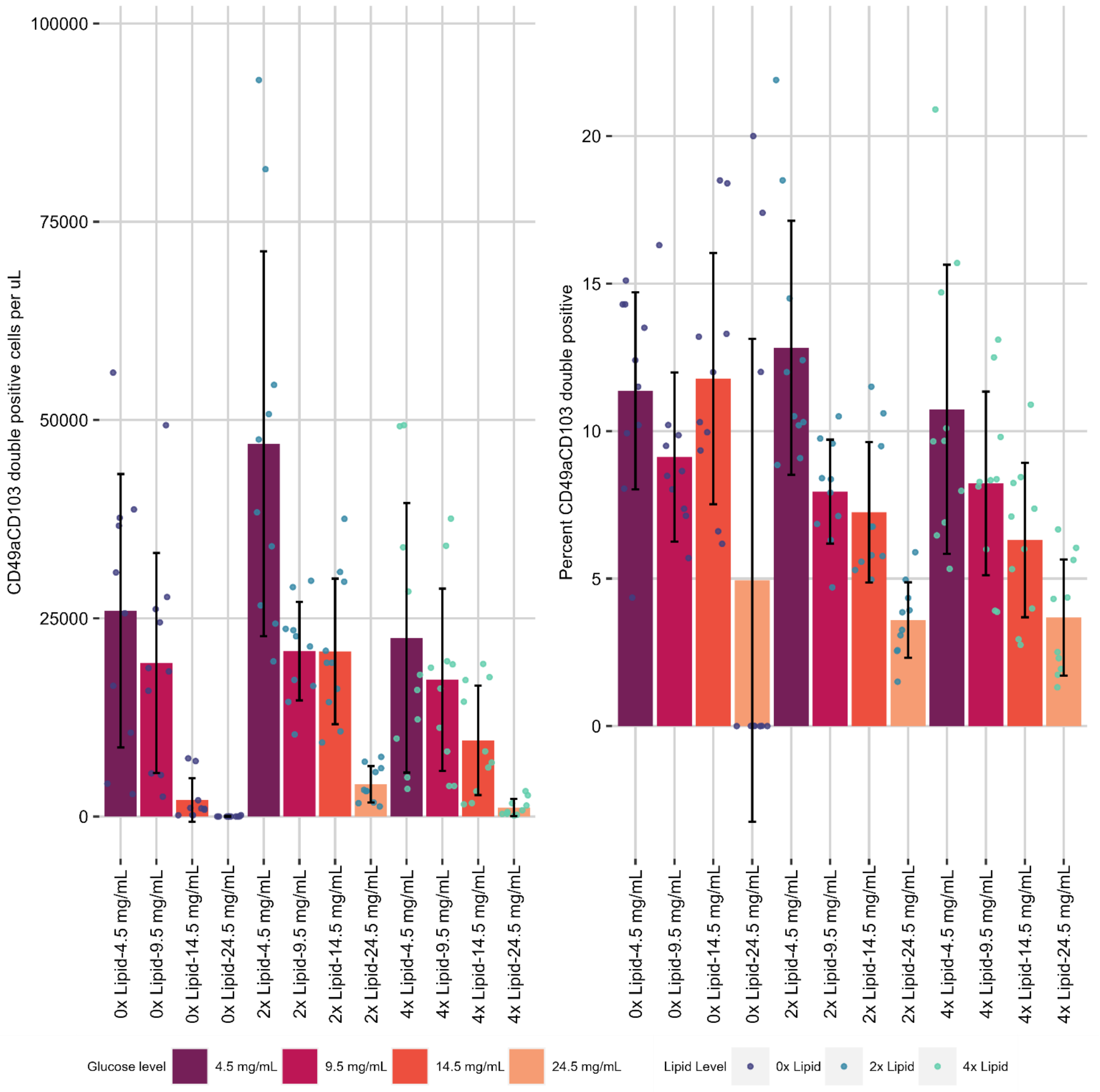


**Supplementary Figure 5. Metabolite concentration impact on CD49a$^{pos}$CD103$^{pos}$ T$_{RM}$-like cell differentiation and maintenance *in vitro:*** Percent CD49a$^{pos}$CD103$^{pos}$ (B) and cell number per µL (A) in metabolite alteration assays where both lipid (0x-4x) and glucose (4.5-24.5 mg/mL) were altered. *P < 0.05, **P < 0.01, ***P < 0.001, ****P < 0.0001. Dots represent individual mice. Data were generated from two independent experiments of 5 mice each.

| Marker | Fluorophore | Company | Catalog No. |
|---|---|---|---|
| **Live-dead** | Aqua | Invitrogen | L34966 |
| **Intravenous CD45** | BV785 | Biolegend | 103149 |
| **CD8α** | APC | Biolegend | 100712 |
| **CD8α** | APC-eFluor 780 | Invitrogen | 47-0081-82 |
| **TCRβ** | BV650 | Biolegend | 109251 |
| **CD103** | BV421 | Biolegend | 121422 |
| **CD49a** | PE | BD Biosciences | 562115 |
| **CD49a** | BV711 | BD Biosciences | 564863 |
| **CD44** | APC | Biolegend | 103012 |
| **CCR7** | PE-Dazzle 594 | Biolegend | 120122 |
| **Fc Block: anti-mouse CD16/32** | None | Biolegend | 101302 |

**Supplementary Table 1: Antibody information and catalog numbers.** Additional information to ensure reproducibility in methods.
